# Quantitative Analysis of Global Protein Stability Rates in Tissues

**DOI:** 10.1101/520494

**Authors:** Daniel B. McClatchy, Yu Gao, Mathieu Lavallée-Adam, John R. Yates

## Abstract

Protein degradation is an essential mechanism for maintaining homeostasis in response to internal and external perturbations. Disruption of this process is implicated in many human diseases, but quantitation of global stability rates has not yet been achieved in tissues. We have developed QUAD (Quantification of Azidohomoalanine Degradation), a technique to quantitate global protein degradation using mass spectrometry. Azidohomoalanine (AHA) is pulsed into mouse tissues through their diet. The mice are then returned to a normal diet and the decrease of AHA abundance can be quantitated in the proteome. QUAD analysis reveals that protein stability varied within tissues, but discernible trends in the data suggest that cellular environment is a major factor dictating stability. Within a tissue, different organelles, post-translation modifications, and protein functions were enriched with different stability patterns. Surprisingly, subunits of the TRIC molecular chaperonin possessed markedly distinct stability trajectories in the brain. Further investigation revealed that these subunits also possessed different subcellular localization and expression patterns that were uniquely altered with age and in Alzheimer’s disease transgenic mice, indicating a potential non-canonical chaperonin. Finally, QUAD analysis demonstrated that protein stability is enhanced with age in the brain but not in the liver. Overall, QUAD allows the first global quantitation of protein stability rates in tissues, which may lead to new insights and hypotheses in basic and translational research.

**Summary:** Protein degradation is an important component of the proteostasis network, but no techniques are available to globally quantitate degradation rates in tissues. In this study, we demonstrate a new method QUAD (Quantification of Azidohomoalanine Degradation) that can accurately quantitate degradation rates in tissues. QUAD analysis of mouse tissues reveal that unique degradation trends can define different tissue proteomes. Within a tissue, specific protein characteristics are correlated with different levels of protein stability. Further investigation of the TRIC chaperonin with strikingly different subunit stabilities suggests a non-canonical chaperonin in brain tissue. Consistent with the theory that the proteostasis network is compromised with age, we discovered that protein stability is globally enhanced in brains of old mice compared to young mice.

## Introduction

Proteostasis is the coordinated regulation of many cellular processes, including protein synthesis, degradation, and folding, to maintain a fully functional proteome in response to cellular perturbations. Dysfunction in any of these cellular processes can disrupt the proteome and trigger disease. Due to the abundance of quantitative techniques available to globally measure changes in protein synthesis, it has been the most studied component of proteostasis. For example, ribosomal profiling allows the quantitation of changes in translation efficiency of thousands of mRNAs in almost any biological sample[1]. Likewise, techniques have been developed to quantitate newly synthesized proteins in a discrete time frame[2, 3]. Identification of changes in protein synthesis rates can precede detectable changes in the whole proteome, and possibly portend a disease phenotype. Interventions at early timepoints in a disease have the greatest potential to prevent permanent damage to cells and tissues and early perturbations in the proteome may be a good indicator something is wrong.

Similarly, changes in protein degradation rates can precede detectable changes in whole proteome. Protein degradation, like synthesis, is required to maintain the optimal protein concentration in response to changes in the cellular environment. In addition, proteins need to be continually degraded to prevent the accumulation of damaged proteins (i.e. oxidation, nitrosylation, or mutations) which may cause protein dysfunction or aggregation[4]. There are two main processes that regulate protein stability in a cell. Autophagy is the cellular process in which misfolded proteins and damaged organelles are engulfed by autophagic vesicles (AV) and then fuse with lysosomes, which degrade the AV contents. Alternatively, post-translational ubiquitination targets proteins to be degraded by the multi-subunit protein complex proteasome. Both lysosome and proteasome pathways are tightly regulated, and dysfunction has been linked to various human diseases[5, 6]. For example, autophagy dysfunction has been described as an early event in Alzheimer’s disease (AD), and drugs manipulating autophagy can improve cognitive deficits in AD transgenic mice[7–9]. In addition, the neurodevelopmental cognitive disorder Angelman Syndrome is caused by a mutation in a ubiquitin ligase, which is essential for ubiquitination of specific proteins[10, 11]. There are many techniques to measure global protein stability rates. The general strategy is a “pulse-chase” experiment, where proteins are labeled or tagged and then quantitated by the loss of protein signal with time. For example, quantitative immunoblots have been used to quantitate stability rates after inhibition of protein synthesis in an epitope tagged yeast strain. [12]. Thousands of stability rates have been reported in cells by expressing mRNA fused to fluorescent tags and measuring protein stability with fluorescence-activated cell sorting or microscopy [13, 14]. Numerous studies have measured protein stability through the use of different heavy stable isotopes to label proteins and mass spectrometry (MS) to quantitate the loss of peptide intensity[15–19]. However, these techniques are restricted to cultured cells and there are very few reports of quantitation of protein stability rates in tissues. In one report, rats were fully labeled with heavy nitrogen(^15^N) through a ^15^N diet, and protein stability was studying after switching the rats to a normal ^14^N diet[20]. After 6 months on the ^14^N diet some proteins were still labeled with ^15^N suggesting that these proteins are very stable or extreme long-lived proteins (ELLP). Protein stability studies using ^15^N labeling ELLP strategy [21] of tissue is limited to the identification of long-lived proteins (i.e. many months). A few other tissue stability studies have reported global protein turnover rates using different approaches [22–25]. However, protein turnover rates encompass both synthesis and catabolism rates of proteins and thus, it is difficult to distinguish whether perturbations in synthesis or stability are responsible for changes in protein turnover rates using this strategy.

We propose to use azidohomoalanine (AHA) to quantitate protein stability rates in tissues using pulse-chase labeling coupled with MS, which will provide better temporal resolution than the ELLP strategy. AHA is a non-canonical amino acid that can be inserted into proteins *in vivo* because it is accepted by the endogenous methionine tRNA. Using click chemistry, AHA containing proteins can be covalently bound to a biotin-alkyne and the AHA containing proteins can then be enriched with neutravidin agarose beads. AHA was originally described in the BONCAT(Biorthogonal Non-canonical Amino acid Tagging) method, which involves labeling cultured cells for short time periods to identify newly synthesized proteins using MS[2]. Using PALM (<u>P</u>ulse <u>A</u>HA <u>L</u>abeling in <u>M</u>ammals), it has recently been demonstrated that AHA can be safely incorporated into the proteomes of mouse tissues through a special diet to identify new synthesized proteins[26]. In this study, we used QUAD (<u>Qu</u>antification of <u>A</u>HA <u>D</u>egradation), a new method to quantify protein stability rates in tissues.

## RESULTS

### AHA pulse-chase strategy to quantitate global protein stability rates in tissues

Figure 1A illustrates the QUAD workflow. Twelve one-month old male C57BL/6 mice were placed on an AHA diet as previously described[26]. After 4 days, three mice were sacrificed and tissues were harvested. This group was designated Day0. The remaining mice were returned to a normal mouse diet for various “chase” times. Three mice were sacrificed after three (Day3), seven (Day7), and fourteen (Day14) days on a normal mouse diet. After tissue homogenization, click chemistry was performed to covalently add a biotin-alkyne to any AHA containing proteins. For sample validation, the four timepoints were analyzed with immunoblot analysis. Protein samples were separated by gel electrophoresis and immunoblots were probed with streptavidin-HRP. Immunoreactivity was observed at all time points, with the most observed at Day0 and the least at Day14 (**Figure 1B**). For MS analysis, samples were labeled with either a light or heavy biotin-alkyne to enable quantification based on the calculation of heavy/light ratios [26]. Day0 samples were labeled with the light biotin-alkyne and all other time points were labeled with the heavy biotin-alkyne. The Day0 samples from different mice were combined to generate one internal standard. After the click reactions, this internal standard was mixed 1:1(wt/wt) with samples from individual mice at different time points. As a baseline measurement, an aliquot of Day0 labeled with light biotin-alkyne was mixed 1:1 (wt/wt) with an aliquot of Day0 labeled with heavy biotin-alkyne. Next, the mixtures were digested with trypsin and AHA peptides were enriched with neutravidin beads. The enriched AHA peptides were eluted from the beads and analyzed by MS. The data analysis output is AHA proteins quantified in four heavy/light mixtures: Day0/Day0, Day3/Day0, Day7/Day0, and Day14/Day0. MS analysis was performed with liver and brain tissue. In total, over 500,000 non-unique AHA peptides were identified representing 6614 unique genes.

**Figure1.**
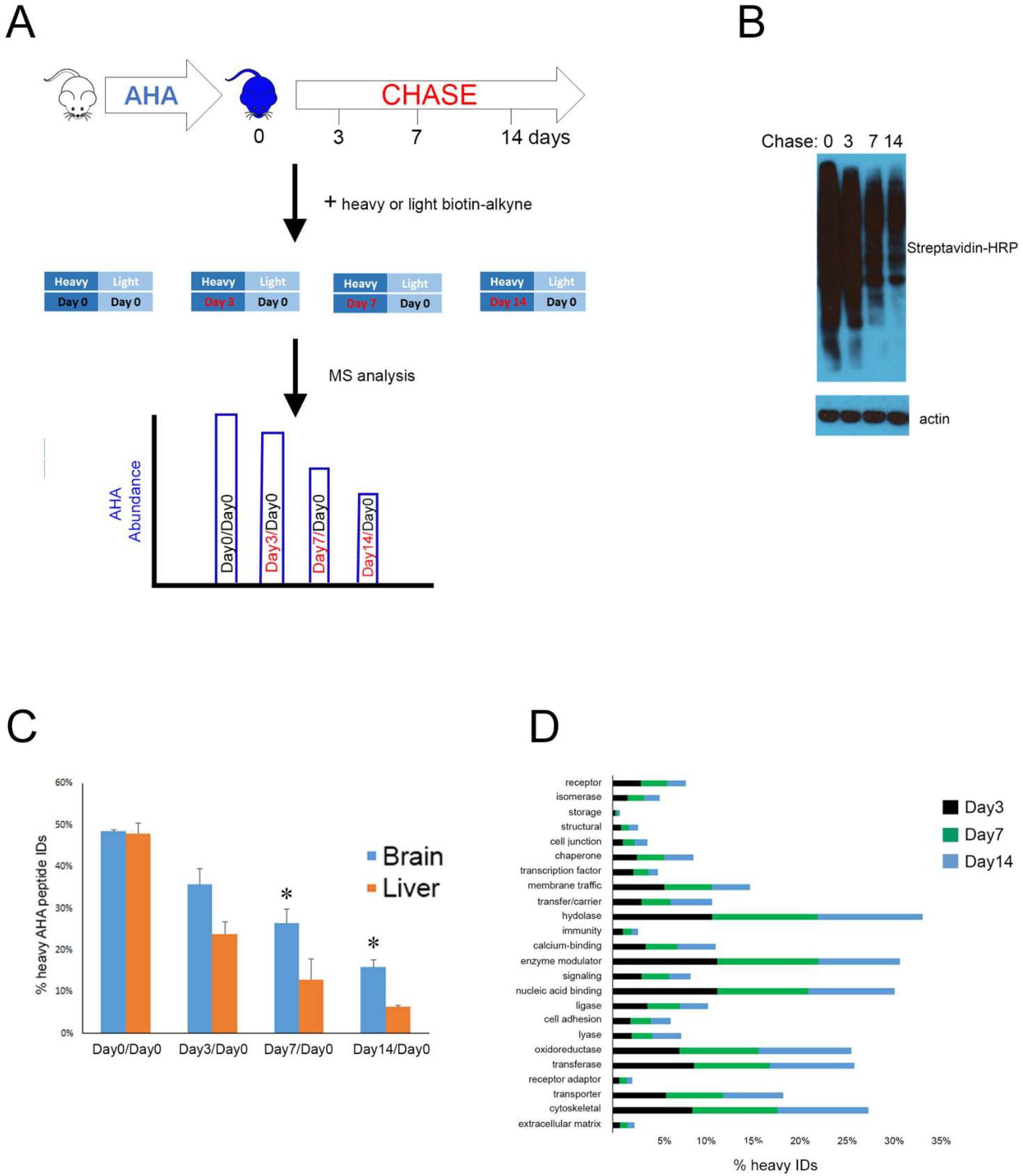
**A**, Schematic of the experimental MS design. **B**, Immunoblot analysis of brain tissue demonstrates a decrease detection of biotin-alkyne with increase chase time. The immunoblot was either probe with streptavidin-HRP or β-actin (loading control). **C**, The number of AHA peptides identified from chase time points decreases with increased chase time. Percentages of heavy AHA peptide identified (y-axis) from the total AHA identifications (i.e. light plus heavy) were calculated from MS analysis of different sample mixtures. The Day0/Day0 consists of two MS analyses of technical replicates and other mixtures represent three MS analyses from three mice. Liver tissue had significantly (* p < 0.05) fewer heavy AHA peptide identifications than brain tissue at Day7 and Day14 using a two-tailed t-test at each chase point. **D**, Identical function protein classes were identified at different chase time points. Y-axis represents the percentage of genes from each dataset (i.e. Day3, Day7, Day14).

First, the percentage of heavy AHA peptides identified in each MS analysis was calculated (**Figure 1C**). For the baseline (i.e. Day0-Heavy/Day0-Light), ~50% of the AHA peptides identified were heavy. With longer chase time, the percentage of heavy AHA peptides identified decreased. Although the baseline was similar between liver and brain, the percentage of heavy AHA peptides identified in the liver was significantly less than in the brain at Day7 and Day14. Since the ability to identify a peptide in the mass spectrometer is directly related to its abundance in the sample, this suggests that heavy AHA proteins become less abundant with longer chase times. The heavy AHA proteins identified at each time point in all three biological replicates were assigned to a large variety of functions, indicating the QUAD strategy is capable of a global analysis of protein stability (**Figure 1D**).

Next, heavy/light ratios were quantified with the algorithm pQUANT[27]. A correlation value (r) of 0.87 between biological replicates suggested good reproducibility, thus allowing for accurate quantification (**Figure 2A**). The median AHA peptide ratio for each protein was calculated and plotted in a histogram (**Figure 2B and C**). As the chase time became longer, the heavy/light protein ratios were progressively smaller for both tissues. For the baseline analyses, the liver and brain tissues had identical medians, which were close to one. The liver median ratio was consistently smaller than the brain median ratio at all chase time points. For example, the median protein ratio in brain at day3 was −0.46, and in liver it was -1.16. Next, an average “protein stability trajectory” or PST was graphed for each AHA protein that was quantified at all chase time points. A wide distribution of trajectories was observed in both tissues, but compared to the brain, more trajectories from the liver had a steeper slope (**Figure 2D and E;TableS1 and S2**). To further investigate, the slope was calculated for each PST. A slope of zero would indicate no change in AHA protein abundance over time (i.e. very stable). The distribution of slopes between tissues overlapped, but a clear trend was apparent. The average brain slope (−0.11) was significantly (p<0.0001) different from liver (−0.16) (**Figure2F**). In the brain, the myelin basic protein(MBP), sirtuin-2, and 2′,3′-cyclic-nucleotide 3′-phosphodiesterase(CNP) were among the most stable proteins while cofilin-1 was one of the least stable(**Figure 2G**), which is consistent with previous reports [20, 23]. Finally, we tested if any intrinsic protein characteristics could contribute to the differences in PST. There was no correlation between PST and molecular weight (**Figure 2H**), abundance (**Figure 2I**), transmembrane regions (**Figure 2J**) or protein disorder (**Figure 3A-C**). Overall, this data demonstrates that protein stability varies greatly within a tissue proteome, but global patterns can distinguish different tissues.

**Figure 2.**
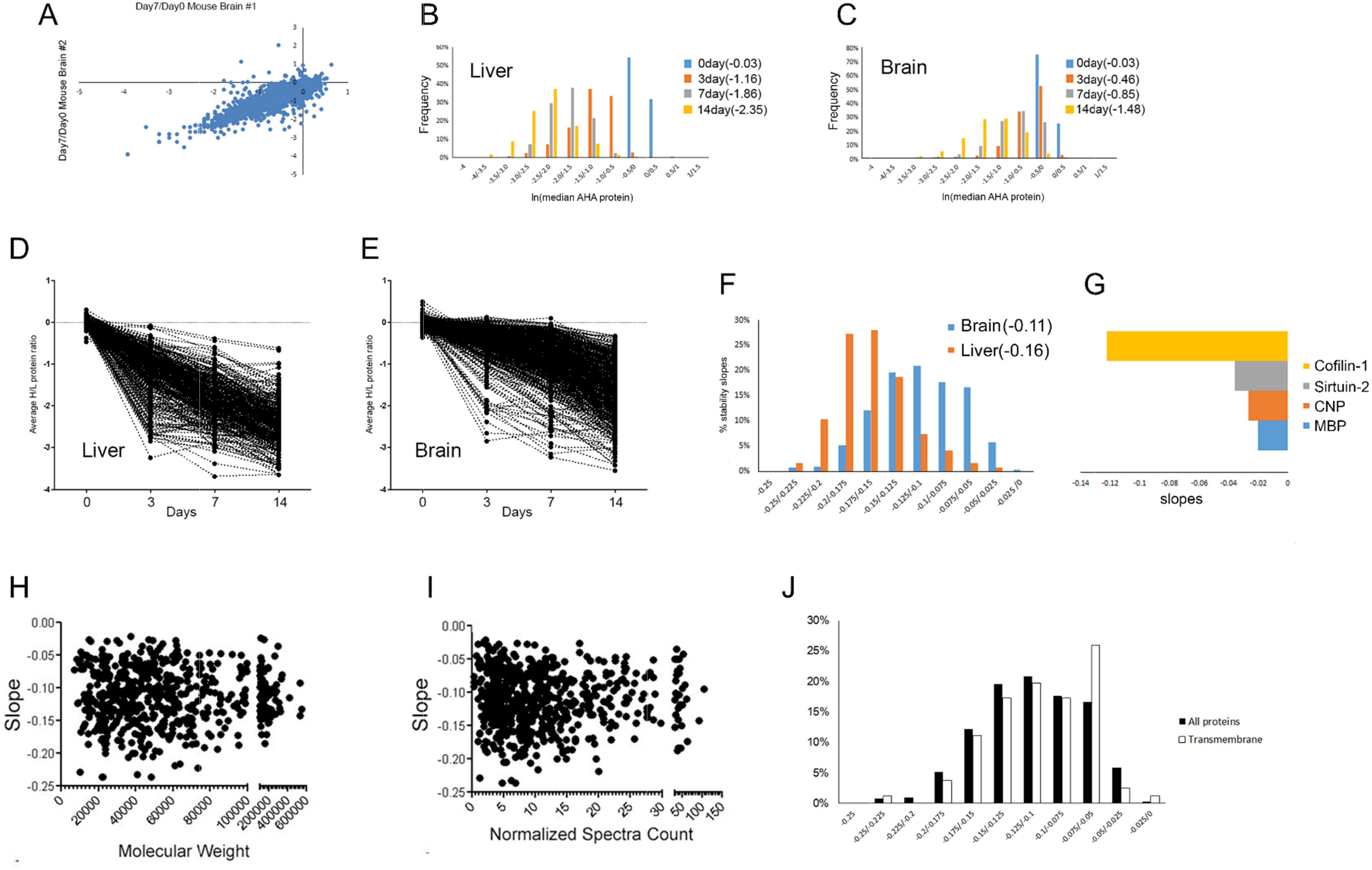
**A**, The correlation value (r) is 0.87 between two biological replicates. The abundance of the AHA proteins decreases with chase time in both liver (**B**) and brain(**C**). The median AHA peptide heavy/light ratios were calculated for each protein at each chase time point. After a natural log transformation, they were plotted in a histogram with the y-axis representing the percentage of proteins. The median protein ratio for each time point is in parentheses in the legend. PST are calculated for liver(**D, TableS2**) and brain(**E, TableS1**). The average AHA protein ratio was calculated for three mice at each chase time point. **F**, The distribution of the slopes of the PST is unique for each tissue. The slopes were calculated for the PST from Figure 2D and 2E and plotted in a histogram with the percentage of slopes on the y-axis. The average slope for each tissue is in parentheses, and was significantly (p<0.0001) different using a two-tail t-test. **G**, Slopes of the PST correspond to previously published reports on protein stability using different methods. Y-axis is the natural log of the slope. There was no correlation with PST in the brain dataset with molecular weight(**H**), protein abundance(**I**), or membrane proteins(**J**).

**Figure 3.**
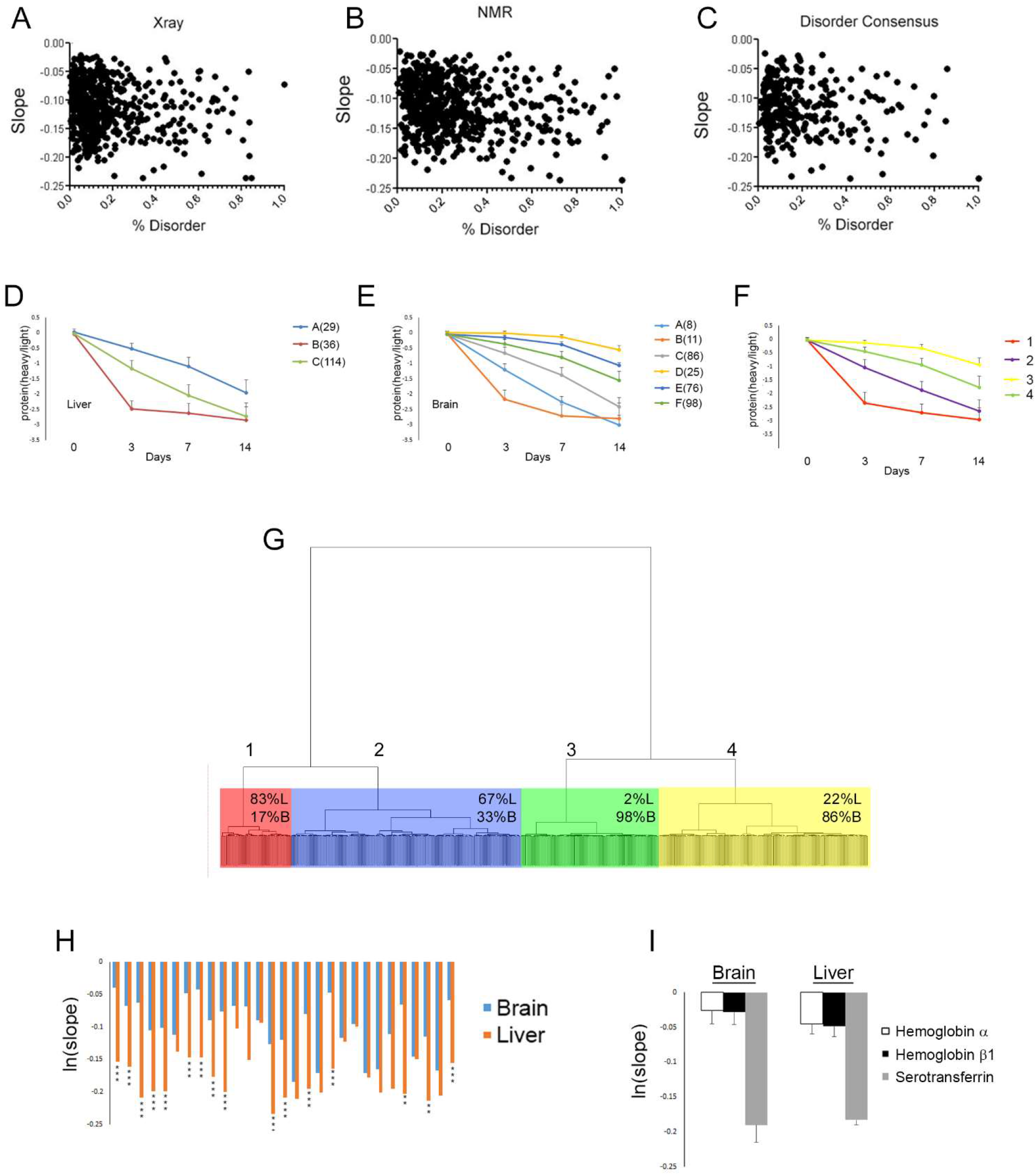
No correlation was observed between PST and predicted protein disorder. Percentage protein disorder predicted(x-axis) from Xray crystallography data (**A**), NMR data(**B**) or a multiple predictor disorder algorithm (**C**) was graphed with the calculated protein slopes(y-axis) from the brain dataset. Clustering analysis was performed on the PST from liver (**D, TableS4**) and brain (**E, TableS3**) separately to identify global trends in each tissue. For each cluster, the average protein heavy/light ratio and standard deviation at each time point was plotted. The number of proteins in each cluster is in parentheses in the legend. **F**, Clustering analysis was performed on PST from liver and brain together. For each cluster, the average protein heavy/light ratio and standard deviation at each time point was plotted(**TableS5**). The number of proteins in each cluster is in parentheses in the legend. **G**, Clusters in F is dominated by one tissue. The dendrogram of the PST summarized in C was annotated with the percentage of trajectories from liver(L) and brain(brain) in each of the clusters. **H**, Statistical analysis of proteins quantified in both liver and brain but assigned to different clusters. Each blue(brain) and orange(liver) pair represent the same protein quantified in both tissues and the y-axis is the natural log transformation of the average slope(**TableS6**). A two-tailed t-test was performed on each protein. **p < 0.01, ***p < 0.001. **I**, Blood and serum proteins had similar quantification results in both tissues. The natural log transformation of the average slope(y-axis) of the average of three biological replicates.

### Cellular environment regulates protein stability

PSTs were further analyzed by an unsupervised learning approach. A clustering analysis was first performed separately on both tissues. Any protein in which the three biological replicate PSTs did not cluster together (meaning they were not similar) was removed from further analysis. Representative clusters were determined based on the average slope and shape trajectory. There were six distinct clusters for brain, and three for liver (**Figure 3D and E;TableS3 and TableS4**). Although some clusters had the same endpoint, the route to that endpoint was different, as illustrated by liver cluster B and C. Clusters with shallow slopes (i.e. cluster D,E, and F) were unique to brain tissue. For a direct comparison, clustering analysis of PSTs was performed on one dataset containing both the liver and brain. Four clusters were clearly distinguishable (**Figure 3F and TableS5**). Proteins from both tissues were present in all clusters, but the clusters were biased toward one tissue (**Figure 3G**). For example, cluster#1 consisted of 83% liver proteins while cluster#3 consisted of only 2% liver proteins. Examining identical proteins that were quantified in both liver and brain tissue but assigned to different clusters, many proteins were significantly less stable in the liver compared to the brain (**Figure 3H and TableS6**). No proteins were observed to be significantly less stable in the brain compared to the liver. Liver and brain proteins annotated to the same clusters were enriched in serum and blood proteins and no differences were observed between the stability of these proteins (**Figure 3I**). Therefore, this analysis indicates that the different cellular environments can affect the stability of the same protein differentially.

### Subcellular localization and protein function can influence protein stability

Next, we investigated whether protein stability is associated with any cellular characteristics within a tissue, such as localization, function, post-translational modifications or protein-protein interactions. For this analysis, the brain dataset in Figure 3E was employed because it contained a wider range of PST than the liver. Clusters D, E, and F were classified as “stable” proteins and clusters A, B, and C were classified as “unstable” proteins. First, GO enrichment analysis of subcellular compartments was performed (**Figure 4A**). Different mitochondrial compartments and the actin cytoskeleton were significantly enriched in the stable dataset, while cytosol, endosomes, axons, and perinuclear compartment were significantly enriched in the unstable dataset. Second, proteins implicated in neurological diseases were analyzed. The entire brain dataset was significantly enriched in proteins reported in Alzheimer’s disease, Huntington’s disease and schizophrenia (**Figure 4B**), but there was no difference in the distribution of stable and unstable proteins associated with these diseases (**Figure 4C and TableS7**). Next, it was determined if a signaling pathway or cellular function was associated with protein stability. The most significant pathway enriched in the unstable dataset (p value = 2.05 e^−16^) was protein metabolism, which included protein synthesis (**Figure 4D and TableS8**). In contrast, the stable dataset was significantly enriched (p value =1.43 e^−16^) in the cytoskeleton, which included cytoskeleton proteins and proteins that regulate the cytoskeleton (**Figure 4E and TableS9**). This functional enrichment was validated with antibodies. Click reactions were performed on brain homogenates from 3 day and 14-day mice. AHA proteins were then enriched with neutravidin beads and the enriched proteins were analyzed by immunoblot analysis (**Figure 4F**). Using antibodies for β-actin and elongation initiation factor 2 alpha (EIF2α), a decrease in immunoreactivity suggesting AHA protein degradation was observed at 14day compared to 3day for both antibodies, but the decrease was significantly larger for EIF2α. β-actin decreased ~30% from 3day to 14day while EIF2α decreased ~75%(**Figure 4G**). When the homogenates were analyzed prior to neutravidin enrichment, no obvious differences were observed with either antibody between 3day and 14day. This data demonstrates that protein stability can be correlated with protein function. Cell death was a significantly enriched pathway in both the stable (p-value =7.56e^−11^) and unstable (p-value – 9.07e^−12^) datasets. Network analysis of these proteins revealed many functional relationships between the stable and unstable proteins (**Figure S1**). For example, manipulation of unstable protein dynamitin expression has been reported to alter the function of stable protein superoxide dismutase −1(SOD1)[28]. Mutants of SOD1 can cause amyotrophic lateral sclerosis and induce cell death and dynamitin expression can affect the ability of SOD1 mutants to induce cell death. Thus, stable and unstable proteins can interact functionally in signaling networks.

**Figure 4.**
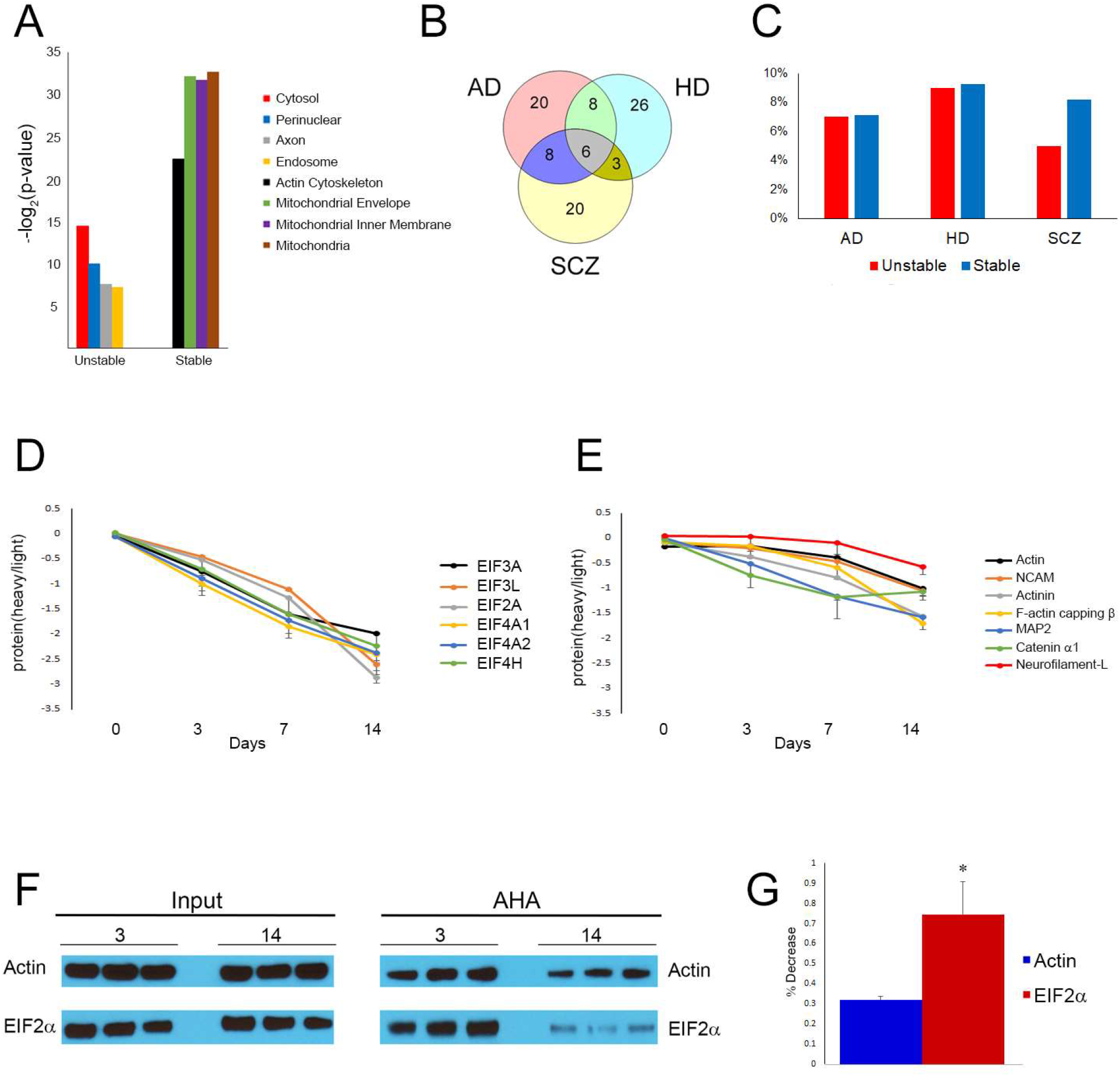
**A**, GO analysis of stable and unstable protein datasets demonstrated significant enrichment of different subcellular components. The four most significant enriched localizations are reported. Y-axis represents the negative log2 of p-value of enrichment. **B**, Proteins were significantly enriched in the pathology of Alzheimer’s disease (AD, p=1.7^−14^), Huntington’s disease (HD, p = 1.5 ^−14^) and Schizophrenia (SCZ, p=3.7 ^−14^). **C**, The percentage of unique disease protein in B that are in the stable or unstable datasets(**TableS7**). Proteins involved in translation are enriched in the unstable protein dataset (**D, TableS8**) and protein involved in the regulation of the cytoskeleton are enriched in the stable protein dataset (**E, TableS9**). The average protein heavy/light ratio from biological replicates at each time point was plotted after a natural log transformation on the y-axis. **F**, Immunoblot analysis confirmed differences between stability between translational(EIF2α) and cytoskeletal(β-actin) proteins. Click reaction was performed on brain homogenates from three mice at 3day and 14day. Samples were analyzed before (Input) and after(AHA) neutravidin enrichment. **G**, Significant difference between the stability of actin and EIF2α was observed with immunoblot analysis. Quantification of the pixel intensity of the immunoreactivity of the enrichment samples in C demonstrated a larger significant (p < 0.05) difference between 3day and 14day with EIF2α than with actin. The y-axis shows the percent decrease in immunoreactivity between 3day and 14day. A two-tailed t-test was performed.

### Post-translational modifications differ between stable and unstable proteins

Post-translation modifications can affect a proteins structure or function. Seven PTMs were examined for their frequency on stable and unstable proteins. For all PTMs the number of modified proteins was similar in both stable and unstable proteins except succinylation (**Figure 5A**). Only 6% of the unstable proteins had reported succinylation sites while 23% were modified in the stable dataset. Next, the number of PTMs per protein was examined (**Figure 5B – H; TableS10**). A significant difference between acetylation, disulfide bonds, and succinylation was observed. The frequencies of acetylation and succinylation were greater in the stable datasets while the frequency of disulfide bonds was greater in the unstable datasets. Overall, this analysis suggests that PTMs can contribute to protein stability.

**Figure 5.**
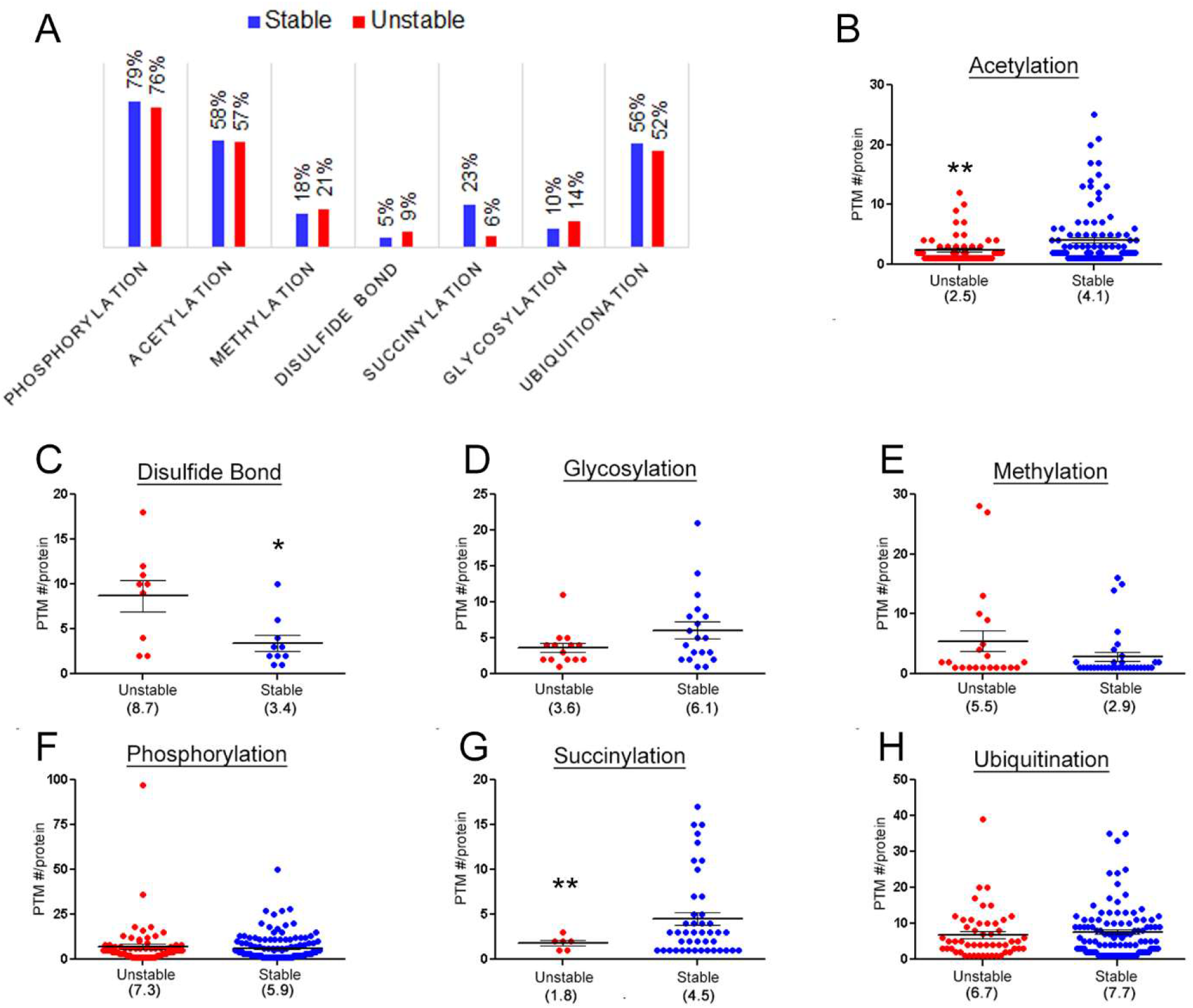
**A**, The percentage of modified proteins in the stable and unstable datasets. The average number of PTM per protein for acetylation (**B**), disulfide bonds (**C**), glycosylation(**D**), methylation (**E**), phosphorylation (**F**), succinylation (**G**), and ubiquitination (**H**). * p < 0.05, ** p < 0.01. **TableS10**

### Protein complexes subunits share similar protein stability

To investigate whether unstable and stable proteins physically interact in *vivo*, the stable and unstable datasets were analyzed together using the STRING interaction database. This analysis was performed conservatively with the highest confidence filter to identify multi-subunit protein complexes opposed to transient direct or indirect (i.e. functional) interacting proteins[29]. The protein complexes identified were the mitochondrial cytochrome C oxidase, mitochondrial NADH dehydrogenase (Complex I), the proteasome complex, and the TRIC (TCP-1 Ring Complex or complex chaperonin containing TCP1 complex (CCT) (**Figure 6A**). All the subunits of both mitochondrial complexes were classified as stable, consistent with the previous localization analysis. The proteasome and the CCT complex sparked further investigation because they contained both stable and unstable subunits. The proteasome is a very large protein structure with over forty subunits that can be divided into the 20S core particle (CP) and the 19S regulatory particle (RP). The CP subunit, Psmb7, was significantly more stable than the two quantified RP subunits, Pmsc1 and Psmd4 (**Figure 6B**). This difference may correspond to reports that the CP and RP are assembled independently [30, 31]. There was also a significant difference observed between two RP subunits, Pmsc1 and Psmd4. Although both are subunits of the RP, they possess different functions. Pmsc1, also known as RPT2, is an ATPase involved in guiding the unfolding and translocation of the ubiquitinated protein into the CP, while Psmd4, also known RPN10, is a non-ATPase subunit involved in the recognition of the ubiquitinated side chain[32, 33]. Thus, differences in protein stability of proteasomal subunits may reflect these previously reported functional differences Differential protein stability with the TRIC chaperonin complex suggests a non-canonical chaperonin in brain tissue.

**Figure 6.**
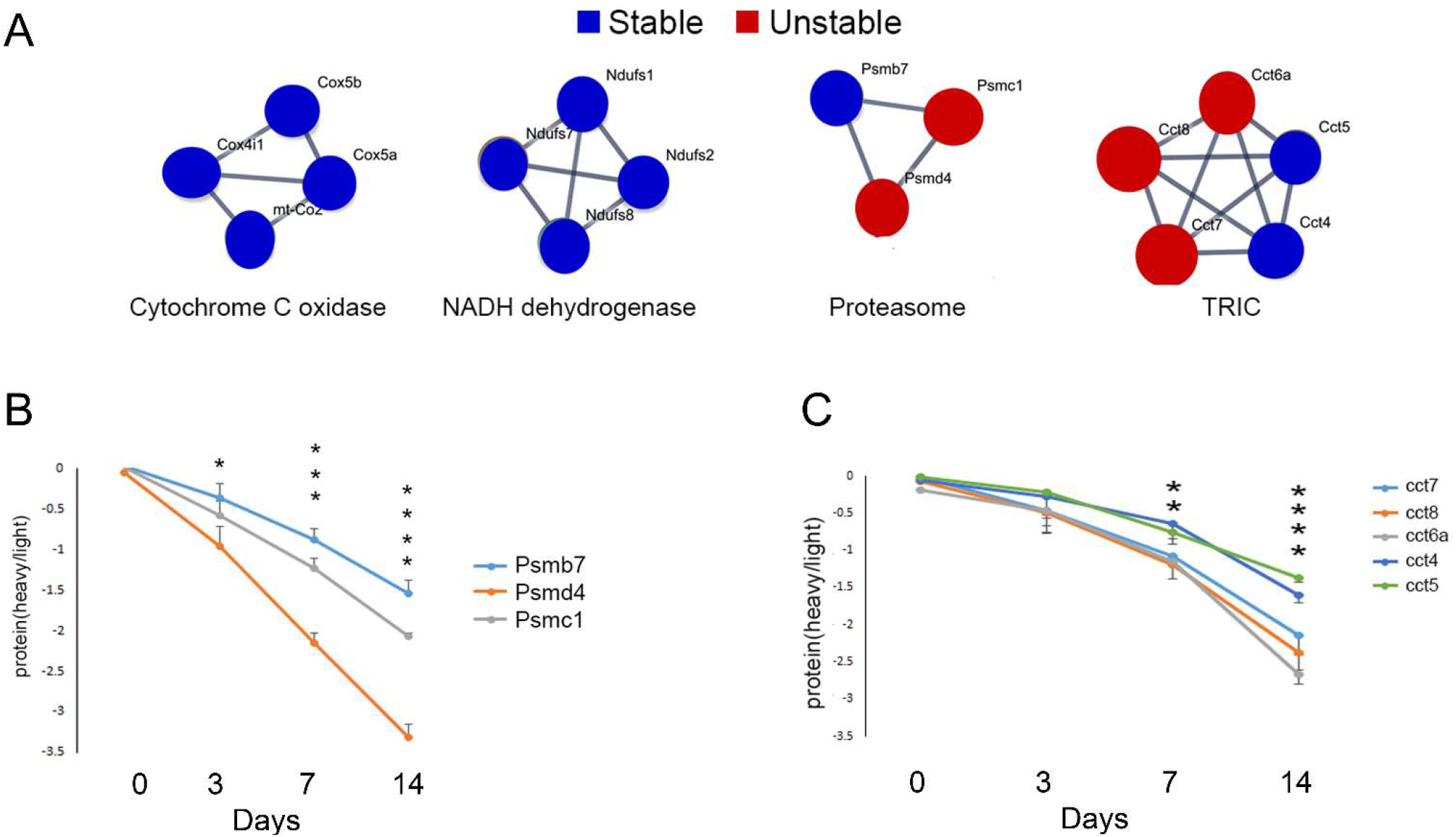
**A**, Protein-protein interactions identified by the STRING Database between stable and unstable proteins. The stability of subunits from four proteins complexes (cytochrome C oxidase, NADH dehydrogenase, proteasome, and TRIC complex) were quantified. Each subunit is annotated with its gene name. **B**, Significant differences in the stability of different proteasome subunits were observed. One-way ANOVA analysis was performed on the subunit protein ratios at each time point. There was a significant difference observed at 3day(p<0.05), 7day(p <0.0001) and 14day(p <0.0001). Bonferroni’s Multiple Comparison post-hoc test was then performed to determine which subunits were significantly different. At 3day, Psmd4 vs Psmb7*. At 7day, Psmd4 vs Psmb7**** and Psmd4 vs Psmc1***. At 14day, Psmd4 vs Psmb7****, Psmd4 vs Psmc1**** and Psmb7 vs Psmc1**. **C**, Significant differences in the stability of different TRIC complex subunits were observed in brain tissue. The average protein heavy/light ratio from biological replicates at each time point was plotted for subunits of the TRiC complex from brain tissue. One-way ANOVA analysis was performed on the subunit protein ratios at each time point. There was a significant difference observed at 7day(p <0.01) and 14day(p <0.0001). Bonferroni’s Multiple Comparison post-hoc test was then performed to determine which subunits were significantly different. At 7day, CCT4 vs CCT6a* and CCT4 vs CCT8*. At 14day, CCT4 vs CCT6a***,CCT4 vs CCT7*, and CCT4 vs CCT8**, CCT5 vs CCT6a****, CCT5 vs CCT7**, and CCT5 vs CCT8***. Figures depict the ANOVA p-values. *p<0.05, **p<0.01, ***p<0.001,****p<0.0001.

The TRiC complex is a molecular chaperonin that consists of two ring structures with each ring comprising eight subunits (CCT 1-8). The subunits are structurally similar, with an ATP-binding equatorial domain and an apical substrate-binding domain linked together by an intermediate domain[34]. The CCT6a, CCT7, and CCT8 subunits clustered in stable brain cluster C, while CCT4 and CCT5 appeared in the unstable brain Cluster F. Statistical analysis confirmed that CCT6a, CCT7, and CCT8 are significantly less stable than the CCT4 and CCT5 subunits (**Figure 6C**). Unlike the proteasomal subunits, stability differences could not be reconciled with reports of CCT subunits in the literature. Since the TRIC complex structure has not been studied in brain, we postulated that a non-canonical TRIC complex may exist in the brain. To provide additional evidence for non-canonical TRIC complexes, experiments were performed to determine if localization differences exist between stable and unstable CCT subunits. Commercial CCT antibodies failed to produce specific staining for immunohistochemistry, so sucrose fractionation was instead employed to examine the nuclear, synaptosomal, and mitochondrial fractions[35]. CCT subunits can be found in published proteomic datasets of these fractions[36–38]. We used immunoblots to compare the immunoreactivity (IR) of CCT5 and CCT8 in total brain homogenate to these fractions, and found that CCT5 was significantly enriched in the synaptosomal and nuclear compartments compared to CCT8 (**Figure 7A-C**). The CCT subunits were detected in the mitochondrial fraction but in much lower abundance than the other fractions. Detection of mitochondrial CCT IR led to saturation of the CCT IR in the total homogenate, preventing quantitation of the immunoblots. The TRIC complex is a component of the proteostasis network, which has been demonstrated to be modified with age [39]. Therefore, we examined the effect of age on CCT expression. There was a significant increase in CCT5 expression in 1-year old mouse brain compared to 2-month old, but no detectable change in CCT8 expression (**Figure 7D-F**). The CCT5 expression change was restricted to the hippocampus (HIP) and cortex (CTX) which corresponded to a significant increase in the synaptosomal and mitochondrial fractions (**Figure 7G**). The HIP and CTX are vulnerable brain regions to AD pathology, and age is a major risk factor for AD. Although how aging contributes to AD pathology is unclear, it has been hypothesized that aging-induced changes in the proteostasis network are involved[39]. Next, CCT expression in an AD transgenic mouse model was examined. A downregulation of both CCT5 and CCT8 in the mitochondrial fraction in 1-year old AD transgenic mouse brains compared to WT littermates was observed (**Figure 7H**). In AD mouse brains before pathology (i.e. 2 months), a downregulation of CCT5 and CCT8 in the mitochondrial fraction was also observed (**Figure 7I**). Interestingly, there was an increase in total homogenate in AD compared to WT, suggesting that CCT5 and CCT8 trafficking in or out of the mitochondrial fraction to the cytosol could be misregulated. Overall, protein stability differences in the subunits of TRIC complex led to the identification of other biological differences between the subunits.

**Figure 7.**
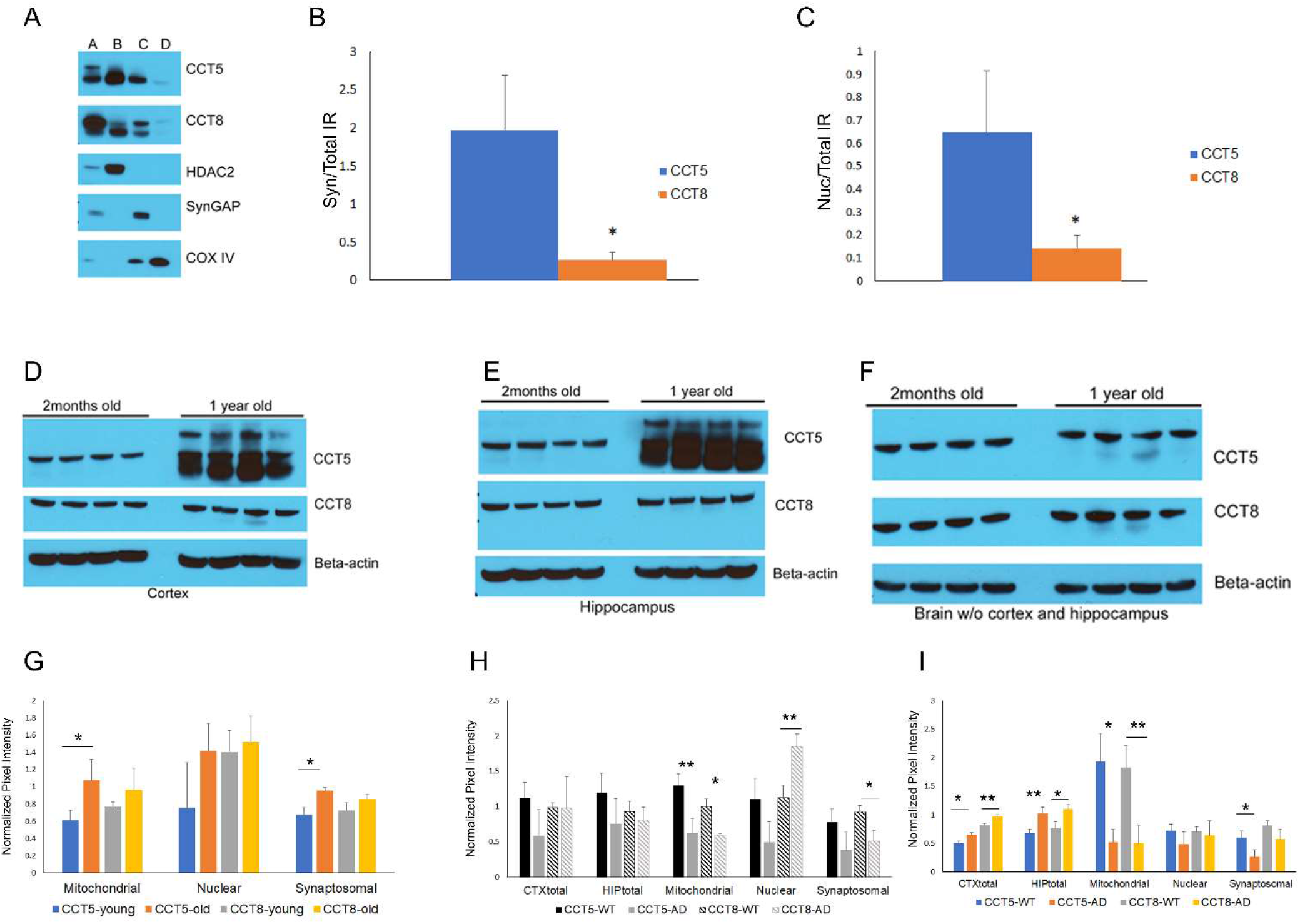
**A**, Representative immunoblot of fractionated brain tissue. A = unfractionated brain, B = nuclear fraction, C= synaptosomal fraction, and D = nuclear fraction. **B**, Quantitation of the CCT5 and 8 IR in the synaptosomal (**B**) and nuclear (**C**) fraction normalized to the CCT IR in unfractionated brain; N=4. Unfractionated CTX(**D**), HIP(**E**) and brain without the CTX and HIP (**F**) from 2month and 1-year old mice probed with CCT5, CCT8, and beta-actin (i.e. loading control) antibodies; N=4. **G**, Quantitation of immunoblots of fractionated CTX/HIP from 2month and 1-year old mice probed with CCT antibodies; N=3. **H**, Quantitation of immunoblots of fractionated CTX/HIP from WT and AD 1-year old mice probed with CCT antibodies;N=3. **I**, Quantitation of immunoblots of fractionated CTX/HIP from WT and AD 2-month old mice probed with CCT antibodies; N=3. Y-axis is the average pixel intensity of the CCT antibodies normalized with antibodies to subcellular markers. *p<0.05, **p<0.01.

### Age influences protein stability in brain tissue

Since proteostasis has been postulated to decline with age, QUAD analysis was applied to brain and liver tissue from 2-month and 1-year old mice. The mice were labeled with AHA for four days. The mice were either sacrificed after the AHA labeling (Day0) or returned to normal food for seven days (Day7) before sacrifice. For each age group, the Day0 tissues were combined for an internal standard and Day7 tissue from three mice were analyzed separately. Quantitation of the Day7/Day0 ratios were performed with heavy and light biotin-alkynes as previously described. There was a significantly higher percentage of heavy Day7 peptides identified in the 1-year old brains compared to the 2-month old brains but not in liver (**Figure 8A**). There was a higher percentage of heavy Day7 peptides in brain compared liver regardless of age consistent with Figure 1C. This suggests that AHA proteins are more abundant after seven days of normal diet in the 1-year mouse brains compared to 2-month old mouse brains, but age had no effect on the AHA proteins in the liver proteome. Next, only the AHA proteins quantified in three mice at both timepoints were compared. The Day7/Day0 protein ratios were equal or greater in the 1-year old brains compared to the 2-month old brains (**Figure 8B**) but evenly distributed in liver (**Figure 8C**). Thus, this QUAD analysis demonstrates that protein degradation is slower in old brain tissue compared to young but age has no detectable effect on protein stability in liver tissue.

**Figure 8.**
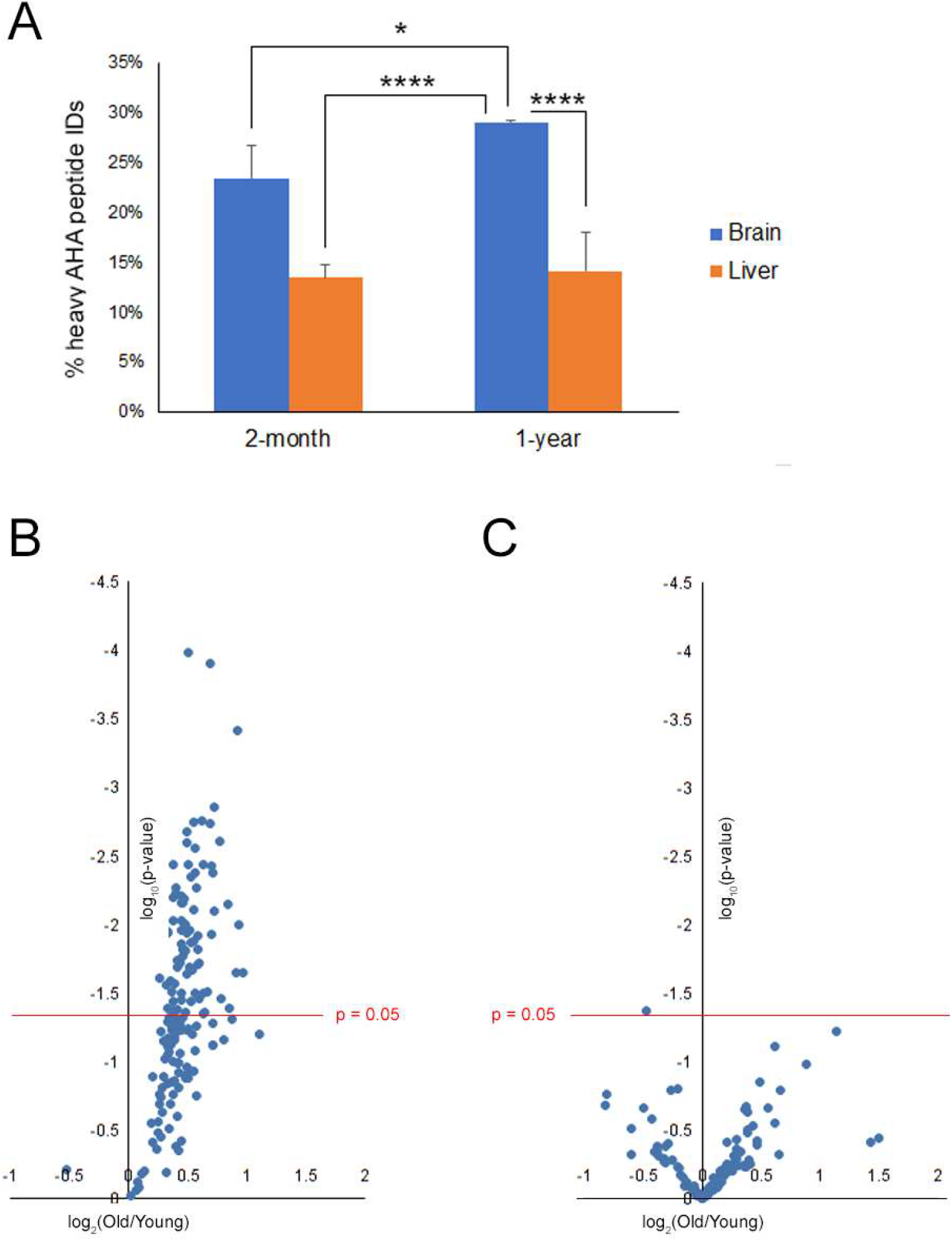
**A**, The number of AHA peptides identified from a 7 day chase period was decreased in 2-month old brains compared to 1-year old brain. Average percentages of heavy AHA peptide identified (y-axis) from the total AHA identifications (i.e. light plus heavy) were calculated from MS analysis. N=3. One-ANOVA analysis with Bonferroni’s post-hoc test was performed. Figure depicts the Bonferroni p-values for 1-year old brain. *p<0.05, **p<0.01, ***p<0.001,****p<0.0001. Bonferroni p-values not in the figure: 2mo-Brain vs. 2mo-Liver **, 2mo-Brain vs. 1yr-Liver **, and 2mo-Liver vs. 1yr-Liver not significant. **B**, AHA proteins have a slower degradation rate in brains from 1-year old mice than 2-month old mice(**TableS11**). **C**, AHA proteins have similar degradation rate in livers from 1-year old mice than 2-month old mice. x-axis is the log2 fold change plotted as 1-year/2-month(**TableS12**). Each point represents the average Day7/Day0 protein ratio, which represents three mice from each age group. The y-axis is the log10 p-value with the red line representing the significant value filter 0.05.

## Discussion

The QUAD method quantitates the loss of AHA from tissue proteomes to accurately quantitate the stability of individual proteins. While previous publications have reported global protein turnover rates in tissues as a proxy for measuring protein stability, changes in protein turnover can results from changes in either protein synthesis, degradation, or both[25]. In addition, tissues have different AA incorporation rates which can interfere with comparisons between tissues[21]. The QUAD method discards the variability of AA incorporation and protein synthesis and solely measures the degradation rates in tissues. Nevertheless, our analysis using QUAD and the protein turnover study by Price et al have both demonstrated different trends between tissues. For example, Price et al. reported that the average protein turnover in liver was faster than in brain, similar to the findings of this study[23]. This is very strong evidence that the cellular environment is one of the main factors that determines protein stability. Our data indicates that measuring degradation alone can define the uniqueness of tissue proteomes.

Within a tissue, there is also a large range of protein stabilities, and stability is not correlated with intrinsic protein characteristics, such as abundance, size, disorder, or structure (i.e. transmembrane regions). However, specific protein localizations and functions could be distinguished by protein stability. Compared to the rest of the proteome, mitochondrial proteins were observed to be more stable, both in this study and in protein turnover studies[23]. It is unclear if this is related to the mitochondrial function or the organelle microenvironment. Translational machinery was enriched in unstable proteins, and cytoskeletal proteins were enriched in stable proteins. Since the cytoskeleton provides the basis for cell polarity and intracellular transport, stability may be needed for these essential structural characteristics. Translation is just as essential for cellular function as the cytoskeleton, but stability of translation machinery may be deleterious to the cell since the overproduction of translational initiation factors is observed in a wide range of cancers[40]. In animal models, it has also been reported that overexpression of initiation factors can increase the susceptibility of tumors, and reduction can suppress tumor development [41–44]. This suggests that tight control of translation factors through degradation is a crucial mechanism to prevent tumorigenesis.

We observed differences in the number of reversible PTMs between stable and unstable proteins. Most PTM research focuses on the function of a single PTM site, so the consequence of significantly more PTM sites on a protein is unclear. It has been postulated that multiple PTMs work in concert to regulate protein function, or possibly, its stability[45]. Disulfide bond frequency was increased in the unstable dataset. Disulfide bonds have traditionally been reported to be required for structural stabilization of nascent proteins, but some reports have demonstrated that this PTM can contribute both positively and negatively to protein stability[46, 47]. It also has been reported to modify protein function and localization[48]. In this study, the unstable proteins with the most disulfide bonds were all secreted or plasma membrane proteins. This is consistent with reports that compared to cytosolic proteins, these protein groups are prone to disulfide bond formation to maintain its structure in the harsh oxidizing extracellular environment[49]. If disulfide bonds do provide protein stabilization, the observation of increased disulfide bonds in unstable proteins seems counterintuitive. One possible explanation is that while disulfide bonds do structurally protect proteins for the extracellular environment, these proteins may still be less stable than proteins that reside in an exclusively intracellular environment. Unlike disulfide bonds, there was an increase number of reported acetylation and succinylation sites on stable proteins compared to unstable proteins, but the exact role and regulation of these PTMs is still under investigation. Global acetylation can be altered by changes in diet, suggesting this PTM may be important for responding to metabolic cues[50]. Investigation into sirtuins, which possess deacetylation activity, has indicated that acetylation regulates the generation of reactive oxygen species(ROS) to provide protection from oxidative damage by modified enzymatic function[51]. For example, sirtuin-3 deacetylates superoxide dismutase 2, which increases its activity to break down ROS[52]. Similarly, dysregulation of succinylation leads to mitochondrial dysfunction[53]. Global unbiased analysis has demonstrated that both acetylation and succinylation are enriched on mitochondrial proteins and there is a large overlap between these PTM sites[53, 54]. There are also reports that acetylation can regulate protein stability. For instance, acetylation has been reported to increase the stability of the toxic cleavage product of ataxin-7[55], but it has also been reported to decrease stability of huntingtin[56]. Thus, regulation of protein stability by acetylation may be context specific. Further investigation is required to determine if the increases in acetylation and succinylation frequency contribute to the enhance protein stability observed in the mitochondria proteome.

We also detected significant stability differences between protein subunits of the TRIC chaperonin complex and postulated that this might indicate a non-canonical chaperonin in the brain. The identification of subunit differences in subcellular localization and age-related expression provides further evidence for this hypothesis. It has been reported that all subunits are required for chaperonin function, as deletion or mutation of any subunits is sufficient to impair the function of the chaperonin in cultured cells [57]. Detailed structure analyses have confirmed the existence of the eight-subunit chaperonin[58–60], but there has been indirect evidence to suggest an alternative structure. One of the first pieces of evidence was a large difference in the subunit mRNA levels in mouse testes[61]. In cultured cells, exogenously expressed subunits have been localized to different subcellular compartments[62, 63]. For example, the application of interleukin-6 causes exogenous CCT3, but not exogenous CCT7, to translocate from the cytoplasm to the nucleus[64]. Furthermore, numerous reports have demonstrated that exogenous expression of subunits reveals subunit specific phenotypes in cultured cells[62, 65–67]. The CCT proteins have not been studied in brain tissue, but the mRNA has. An unequal distribution in different regions was reported in the mouse brain[68]. In human brain, all CCT mRNAs were decreased with age, except CCT5 which was increased in the hippocampus, similar our observations in mouse brain[39]. In addition, compared to unaffected human brains, the CCT5 and CCT8 mRNAs are significantly decreased in human AD brain, but not in Huntington’s or Parkinson’s disease brains. AD is the most common dementia with 5 million Americans affected, and the number of cases is expected to triple by 2050[7]. AD pathology is defined as extracellular plaques consisting of amyloid beta peptide (Aβ), neurofibrillary tangles consisting of the hyperphosphorylated tau protein, and neurodegeneration of the CTX and the HIP, which are the regions responsible for memory and cognition. Aβ is generated from the cleavage of the single transmembrane protein APP (Amyloid Precursor Protein) by β- and γ-secretases. Mutations in APP and the γ-secretase subunits, presenilins (PSEN1 and PSEN2), have been implicated in familial AD. In an AD transgenic mouse model with human mutations in APP and PSEN1, we observed a decrease in both CCT5 and CCT8 in the mitochondrial fraction in 1-year old AD transgenic mice in CTX and HIP. It is unclear if this CCT depression is contributing to or in response to the AD pathology. However, we also observed a decrease in both CCT5 and CCT8 in the mitochondrial fraction in 2-month old AD transgenic mice prior to the detection of AD pathology which suggests that CCT depression may contribute to AD pathology. The TRIC has been localized to the mitochondria associated ER membranes(i.e. MAM)[36]. APP and γ-secretases have also been localized to MAM, and alterations in this subcellular compartment has been proposed to contribute to AD pathogenesis [69]. Our data raises the possibility that the TRIC complex may contribute to Aβ pathology, which has been proposed for other protein folding complexes in neurodegenerative diseases[70]. However, before determining the possible contribution of TRIC to AD pathology, the structure and function of TRIC in different subcellular compartments in the brain needs to be resolved. Overall, this demonstrates how the quantitation of protein stability rates in tissues can lead to new insights and hypotheses in basic and translational research.

Protein degradation is a crucial component of proteostasis, which has been postulated to decline with age. Age-related decline of proteostasis in the brain has received considerable attention because age is a major risk factor for neurodegenerative diseases. A common pathological feature of these disorders is the accumulated and aggregation of misfolded proteins[71]. It has been hypothesized that age-related impairment of protein folding machinery leads to these pathological observations[39]. Our data suggests there is an age-related decline protein degradation. Although this is the first study to quantify global degradation rates in tissues, other laboratories have demonstrated that chemical or genetic modification of autophagy or proteasomal degradation can affect neurological health in old mice. For example, induction of autophagy by rapamycin *in vivo* lowers intracellular amyloid beta levels and improves cognition[8], and long-term rapamycin treatment reduces plaque load in AD mouse models[9]. Thus, we postulate that a global decline in protein degradation in brain tissue contributes to the vulnerability of the elderly to neurological diseases associated with protein misfolding.

In summary, deleterious changes in protein degradation has been implicated in diseases in almost every human tissue. QUAD analysis allows the global quantification of protein stability rates in any mouse tissue which then can be extended to any mouse model of disease. The temporal resolution of QUAD can identify alterations in protein stability prior to development of disease phenotypes, thus identifying potential targets to ameliorate or prevent pathogenesis. With the development of non-canonical amino acids with cell-type specificity [72], AHA can be replaced to allow QUAD analysis to quantitate cell-specific protein stability in animal models of disease.

## Acknowledgements

This work has been supported by NIH grants P41 GM103533, R01 MH067880 (to J.R.Y.). We thank Dr. Claire Delahunty(TSRI) for critical review of the manuscript.

## Author Contributions

Conceptualization, D.B.M and J.R.Y.; Investigation, D.B.M; Formal Analysis and Visualization, D.B.M, Y.G., and M.L.; Writing – Original Draft, D.B.M.; Writing – Review & Editing, D.B.M., Y.G., M.L., and J.R.Y.; Funding Acquisition and Project Administration, J.R.Y.; Supervision, J.R.Y.

## Materials and Methods

### Animals

Mice were housed in plastic cages located inside a temperature- and humidity-controlled animal colony and were maintained on a reversed day/night cycle (lights on from 7:00 P.M. to 7:00 A.M.). Animal facilities were AAALAC-approved, and protocols were in accordance with the IACUC. Male C57BL/6 mice were used for all experiments except for Figure 8 where 2-month and 1-year old mice were used. For the QUAD analysis, mice were fed the AHA diet as previously described for 4 days[26]. AHA was purchased from Click Chemistry Tools (Scottsdale, AZ) and given to Envigo (Madison, WI) to manufacture the AHA mouse pellets. After 4days, the mice were either sacrificed or return to normal mouse feed for various times described in the Results section. The AD transgenic mice used in this study were B6C3-Tg(APPswe,PSEN1dE9)85Dbo/Mmjax– 129x1/SvJ[73]. Only males were used at the ages described. Animals were anesthetized with halothane and sacrificed by decapitation. The whole tissues were quickly removed, dissected, and snap-frozen in liquid nitrogen.

### Tissue Preparation

Thawed liver tissue was further dissected into small pieces and homogenized at 4C using the Precellys 24 homogenizer in PBS with protease and phosphatase inhibitors (Roche, Indianapolis, Indiana). Brain tissue was homogenized in a teflon dounce grinder on ice in PBS with protease and phosphatase inhibitors (Roche, Indianapolis, Indiana). After homogenization, protein concentration was determined with a Pierce BCA protein assay (Life Technologies, Grand Island, NY).

### Click Chemistry

For MS analysis, 10mg were used for each biological replicate plus 10mg for the internal standard(Day0) except for Figure 8 where 5mg were used for each biological replicate plus 5mg for the internal standard(Day0). For immunoblot analysis (Figure 4F), 4mg was used. Sodium dodecyl sulfate was added to the homogenized tissues at final concentration of 0.5%. The homogenate was then sonicated with a tip sonicator and was divided into 0.5 mg aliquots. A click reaction was performed on each aliquot. The click reaction protocol has been previously published[74]. In brief, for each click reaction, the following reagents were added in this order: (1) 30 μL of 1.7mM TBTA, (2) 8 μL of 50 mM copper sulfate, (3) 8 μL of 5mM light biotin-alkyne (C_16_H_24_N_4_O_3_S, Seterah, Eugene, OR) or heavy biotin-alkyne (C_13_H_24_N_3_O_3_S^13^C_3_^15^N, Seterah, Eugene, OR), and (4) 8 μL of 50 mM TCEP. PBS was then added to a final volume of 400 μL and incubated for 1 h at room temperature. The click reactions were combined for each sample and precipitation was performed with 25% TCA.

### Digestion and Biotin Peptide Enrichment

Precipitated pellets were resuspended with MS-compatible surfactant ProteaseMAX (Promega, Madison, WI) and urea, then reduced, alkylated, and digested with TrypZean trypsin (St. Louis, MO, Sigma-Aldrich) at 1:25 dilution with the protein sample as previously described[26]. The digested solution was centrifuged at 13 000g for 10 min. The supernatant was removed, and the pellet was resuspended with PBS and centrifuged at 13 000g for 10 min. Supernatants were combined and 300 μL of neutravidin agarose resin (Thermo Fisher Scientific, Rockland, IL) was added. The resin was incubated with the peptides for 2 h at room temperature while rotating; then, the resin was washed five times with PBS. The peptides were eluted four times with 250 μl 80% acetonitrile, 0.2% formic acid, and 0.1% TFA. The elutions were dried with a speed-vac and stored at −80°C until MS analysis.

### Mass Spectrometry Analysis

Enriched peptides were analyzed by MudPIT. Briefly, Dried peptides were resolubilized in Buffer A (5% ACN, 95% water, 0.1% formic acid) and then were pressure-loaded onto a 250-μm i.d. capillary with a kasil frit. The capillary contained 2cm of 10 μm Jupiter C18-A material (Phenomenex, Ventura, CA), followed by 2 cm 5 μm Partisphere strong cation exchanger (Whatman, Clifton, NJ). This loading column waswashed with buffer A. After washing, a 100 μm i.d. capillary with a 5 μm pulled tip packed with 15 cm 4 μm Jupiter C18 material (Phenomenex, Ventura, CA) was attached to the loading column with a union, and the entire split-column(loading column-union-analytical column) was placed inline with an Agilent 1100 quaternary HPLC (Palo Alto, CA). The sample was analyzed using MudPIT, which is an eleven salt-step separation previously described[75]. As peptides eluted from the microcapillary column, they were electrosprayed directly into a Velos mass spectrometer (ThermoFisher, Palo Alto, CA) with the application of a distal 2.4 kV spray voltage. A cycle of one full-scan FT mass spectrum (300-1600 m/z) at 60 000 resolution followed by 20 data-dependent IT MS/MS spectra at a 35% normalized collision energy was repeated continuously throughout each step of the multidimensional separation. For the analysis in Figure 8, a nano-Easy HPLC (ThermoFisher) with an Elite mass spectrometer (ThermoFisher) was used with the MS setting previously described[75].

### Analysis of Mass spectra

MS1 and MS2 (tandem mass spectra) were extracted from the XCalibur data system format (.RAW) into MS1 and MS2 formats using RawExtract[76]. The MS2 files were interpreted by Prolucid and results were filtered, sorted, and displayed using the DTA Select 2 program using a decoy database strategy filtering for only fully tryptic peptides with a 5ppm mass accuracy [77, 78]. Searches were performed against UniProt mouse database released on 03-25-2014. No enzyme specificity was considered for any search. The following modifications were searched for: static modification of 57.02146 on cysteine for all analyses, and a differential modification of 351.1774 (heavy) or 347.1702 (light) on methionine for AHA bound to a biotin-alkyne.The protein false discovery rate was < 1%. pQuant used the MS1 and DTASelect-filter files for the quantification of the heavy/light ratios using a 0.1 quality filter as previously described[26, 27].

### Sucrose Fractionation of Brain Tissue

Brain tissue was fractionated as previously described[35]. Briefly, whole brains or dissected brain regions were homogenized in 4mM HEPES(pH 7.4), 0.32M sucrose (i.e. Buffer H) using a Teflon dounce grinder. Homogenates were centrifuged at 800 × g at 15 minutes at 4°C. The pellet was resuspended in buffer H and centrifuged at 800 × g at 15 minutes at 4°C. The nuclear pellet was saved and the two supernants were combined. The supernant was then centrifuged at 10, 000 × g for 15 minutes at 4°C. The pellet was resuspended in buffer H and fractionated using a discontinuous sucrose gradient consisting of 0.85, 1.0, and 1.2M sucrose at 100,000 × g for 2 hours at 4°C. After centrifugation, the synaptosomal and mitochondrial fractions were isolated at the 1.0/1.2 interface and the pellet respectively. The nuclear pellet was resuspended in Buffer H with 0.5% NP-40 and incubated on ice for 1hour. The sample was then centrifuged at 1000 x g for 10min. The pellets were washed with Buffer H with 0.5% NP-40 three times. Protein concentration was determined with a Pierce BCA protein assay (Life Technologies, Grand Island, NY).

### Immunoblot Analysis

For Figure 4F: After protein enrichment with neutravidin beads, the washed beads were resuspended using 4X Laemmli Sample Buffer (Bio-Rad) with β-mercaptoethanol. All other immunoblot samples also employed 4X Laemmli Sample Buffer. Samples were separated with 4–12% Bis-Tris gradient gel(Life Technologies), transferred to PVDF blotting paper, and developed as previously described[79]. The immunoblotting antibodies were β-actin(Sigma#A5441), CCT5(Scbt#sc-377261), CCT8(Scbt#sc-376188), COXIV(CST#4850), EIF1A(CST#2538), H3 Histone(ProteinTech#17168-1), HDAC2(ProteinTech#12922-3), Synaptophysin(Abcam#ab8049), and SynGAP(Abcam#ab3344). The immunoblots were quantitated as previously described[35] using different markers to normalize: β-actin(unfractionated), nuclear fraction(H3 Histone), synaptosomal fraction (Synaptophysin) and mitochondrial fraction(COXIV).

### Bioinformatic Analysis

Protein function was assigned using Panther in Figure 1D[80]. Transmembrane proteins (Figure 2J) were determined using UniprotKB[81]. For the disorder correlation (Figure 3A-C), Mobi-lite software was used to determine the Disorder Consensus[82] and Esprite software was used to determine disorder from Xray and NMR databases[83]. PST clustering analysis was performed using Ward’s algorithm[84] and Euclidean distance as a distance measure between PSTs. This clustering analysis was implemented in R for dendrogram generation using the OompaBase and ClassDiscovery packages. Clusters were determined upon dendrogram visual inspection. Localization analysis in Figure 4A was performed by FunCoup[85, 86]. Ingenuity Pathway Analysis was used to calculate significantly enriched diseased associated proteins(Figure 4B and 4C), and cellular functions(Figure 4D,E and F)[87]. The reported PTMs were extracted from UniprotKB except ubiquitination for which the mUbiSiDa database was used[88]. For protein interaction analysis(Figure 5A and B), STRING database was used and an interaction was required to have the highest confidence setting(0.9) and text mining was removed as evidence. All other statistics and graphs were performed in either Microsoft Excel or GraphPad Prism software.

## Supplementary Data

### All raw MS data files will be available on ProteomeXchange after publication

TableS1-All the proteins quantified from brain in all three mice at three timepoints(Day3, Day7, and Day14). Table contains the average heavy/light protein ratio in the mice, relative standard deviation(rsd) of each measurement, and slope of these averages for each protein. All data has been natural log transformed. Data plotted in Figure 2E.

TableS2-All the proteins quantified from liver in all three mice at three timepoints(Day3, Day7, and Day14). Table contains the average heavy/light protein ratio in the mice, relative standard deviation(rsd) of each measurement, and slope of these averages for each protein. All data has been natural log transformed. Data plotted in Figure 2D.

TableS3 – Proteins from brain with their assigned cluster in Figure 3E

TableS4 - Proteins from liver with their assigned cluster in Figure 3D

TableS5 – Proteins from liver and brain with their assigned cluster in Figure 3F and 3G.

TableS6 - The proteins quantified in both liver and brain and the statistical information for Figure 3H.

TableS7-The proteins annotated to be involved in Alzheimer’s disease(AD), Huntington’s disease(HD), and in Schizophrenia(SCZ) in Figure 4B and 4C.

TableS8 – Significantly enriched cellular functions in the unstable brain dataset

TableS9 - Significantly enriched cellular functions in the stable brain dataset

TableS10 – The number of PTM sites report on proteins in our brain dataset in Figure 5.

TableS11-The proteins quantified in Figure8B. The average Day7/Day0 ratios quantified in three 2-month and three 1-year mouse brains.

TableS12-The proteins quantified in Figure8C. The average Day7/Day0 ratios quantified in three 2-month and three 1-year mouse livers.

**FigureS1.**
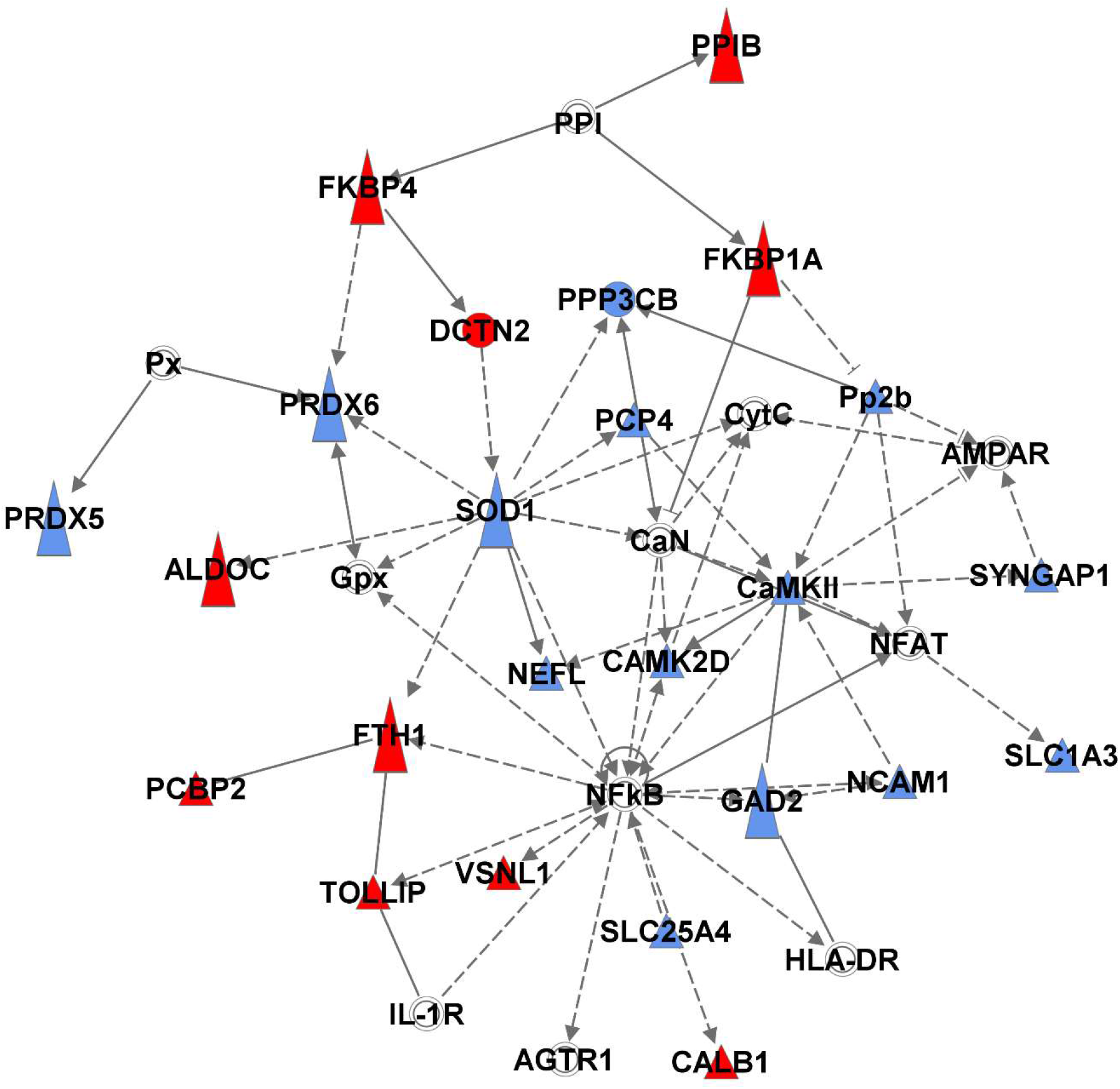
Stable(blue) and unstable(red) proteins form a signaling network that supports cell death. Image was created using Ingenuity. Solid lines represent direct interactions and dotted lines represent indirect interactions. White shapes represent proteins not quantified in the study.

